# Dynamics of mitochondrial mRNA abundance and poly(A) tail during the mammalian oocyte-to-embryo transition

**DOI:** 10.1101/2021.08.29.458072

**Authors:** Yusheng Liu, Yiwei Zhang, Zhonghua Liu, Falong Lu, Jiaqiang Wang

## Abstract

Mitochondria are responsible for producing a cell’s energy and play a central role in many metabolic tasks, as well as signaling transduction and cell death^1^. Mitochondria dysfunctions cause several human diseases and aging processes^2–8^. Mammalian oocytes contain far more mitochondria than somatic cells. The nuclear localization of mitochondrial tricarboxylic acid cycle (TCA) cycle enzymes, which normally localize in the mitochondria, is critical for zygotic genome activation (ZGA) and the oocyte-to-embryo transition (OET) in mice^9^. However, during the mammalian OET, the abundance and post-transcriptional regulation of mitochondrial mRNA (MT-mRNA), particularly the poly(A) tail, has never been studied. Here, we used two independent sequencing methods (PAIso-seq1 and PAIso-seq2) to describe the features of MT-mRNA in mouse cell lines, thirteen mouse tissues and during the OET in mouse, rat, pig, and humans. These features included expression abundance, poly(A) tail length, and non-A residues in poly(A) tails. Unlike nuclear mRNA, we discovered that MT-mRNA has a stable distribution pattern of poly(A) tail length in different cell lines, across tissues, and during mammalian OET. MT-mRNA possesses non-A residues in the poly(A) tail (non-A residues hereafter), which change slightly across tissues and during the OET. We also found that the abundance of MT-mRNA varies substantially across tissues, increases during the OET, and increases along major ZGA in mice, rats, pigs, and humans. These findings provide insights into changes in MT-mRNA abundance and poly(A) tail during the mammalian OET and provide a resource for future studies on the posttranscriptional regulation of mitochondrial transcripts.

## Introduction

Mitochondria are double-membrane-bound organelles found in most eukaryotic organisms. They are known as the “powerhouse of the cell”, since their dominant role is to produce adenosine triphosphate (ATP)^1^. In addition to supplying cellular energy, mitochondria are involved in other cellular processes, including signal transduction, cell growth, cell differentiation, and cell death^1^. Mitochondria are associated with several human diseases, such as cardiac dysfunction^2^, heart failure^3^, autism^4^, diabetes^5^, and Leber Hereditary Optic Neuropathy^6^. Changes to mitochondria also happen during the aging process^7,8^.

Mitochondria play a key role in the development of germ cells and embryos. For example, steroid synthesis occurs in the mitochondria^10^, which is important for gametogenesis. Mitochondria are essential for fertilization, by providing the sperm with energy. Significantly, nuclear localization of mitochondrial tricarboxylic acid cycle (TCA) cycle enzymes is a critical step in mammalian zygotic genome activation (ZGA) and the oocyte-to-embryo transition (OET)^9^.

While most DNA is found in the cell nucleus, mitochondria contain their own genome (mitogenome)^11^. In animals, the mitogenome is typically a single circular chromosome approximately 16 kb long, which has 37 genes^12^. Mitochondrial genes are transcribed as multigenic transcripts, which are cleaved and polyadenylated to generate individual mRNA, tRNAs, and rRNAs^12^. Mitochondria can translate the mitochondria-encoded mRNAs (MT-mRNAs) into 13 protein subunits of the electron transport chain.

Recent developments in deep sequencing have provided a global profile of the nuclear transcriptome. A comprehensive analysis of the human mitochondrial transcriptome across multiple cell lines and tissues has been performed^12^, revealing what has been transcribed from the human mitochondria genome. However, previous transcriptome analysis methods discarded the poly(A) tail information, although most of the MT-mRNAs contains poly(A) tails likely synthesized by mitochondrial poly(A) polymerase^13–17^. Therefore, details on the poly(A) tail of MT-mRNA, including length dynamics or non-A residues, are largely unknown in mitochondria. This is especially true during mammalian oocyte maturation and preimplantation development.

In this study, we provide comprehensive maps of MT-mRNA in cultured cell lines, mice tissues, and oocytes and preimplantation embryos in mice, rats, pigs, and humans via two independent full-length sequencing methods, PAIso-seq1 and PAIso-seq2^18,19^. These analyses reveal the MT-mRNA poly(A) tails are with unique properties and are controlled independent of nuclear mRNA poly(A) tail regulation, providing new insight into the mitochondria function and regulation.

## Results

### Features of MT-mRNA

To identify the distinct features of the MT-mRNA, we first examined PAIso-seq2 datasets of mouse NIH 3T3 cells (3T3) and mouse embryonic stem cells (ES). PAIso-seq2 can read transcriptome-wide full-length RNA isoforms with entire poly(A) tails accurately and reproducibly. While approximately 10% of aligned reads are from mitochondria proteincoding genes (Fig. 1a, b), sufficient coverage is achieved to analyze features of the mitochondrial transcriptome. We provided an expression profile that encompassed all annotated MT-mRNA and found that the expression levels of different genes vary greatly (Fig. 1c). Mitochondrial RNAs are derived from precursor transcripts that traverse almost the entire heavy (Fig. 1a, genes in magenta) and light (Fig. 1a, genes in cyan) mitogenome strands, which are subsequently processed into individual mRNA^12^. As such, the significant variation in levels of MT-mRNA indicates strong post-transcriptional mechanisms, including those affecting transcript stability, in mitochondria. We also found similar expression profiles between 3T3 and ES (Fig. 1c), reflecting conserved posttranscriptional regulation mechanisms in mitochondria.

**Fig. 1.**
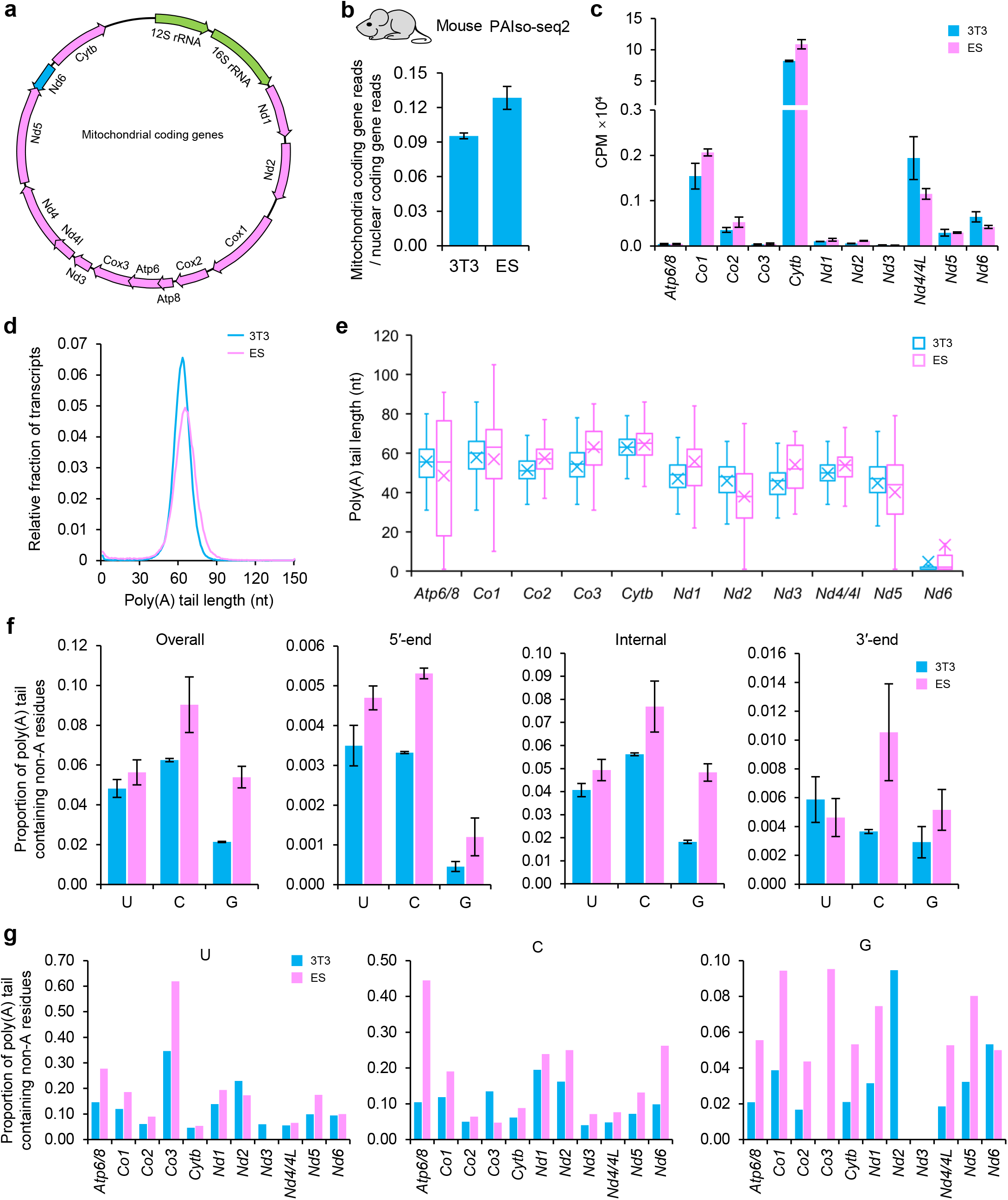
Features of MT-mRNA in 3T3 and mouse embryonic stem (ES) cells. **a,** Map of the mouse mitochondrial genome showing 13 proteins encoded by mitogenomes. Coding genes from heavy chains are shown in red, coding genes from light chains are shown in blue, and rRNAs are shown in green. **b,** MT-mRNA abundance in 3T3 and ES measured by PAIso-seq2. The ratio of reads from mitochondria coding genes to reads from nuclear coding genes is used to present the MT-mRNA abundance. Nuclear coding genes with at least 10 reads in each sample are included in the analysis. Error bars indicate standard error of the mean (SEM) from two replicates. **c,** Expression levels of mitochondrial coding genes in 3T3 and ES measured by PAIso-seq2. Error bars indicate SEM from two replicates. **d,** Histogram of poly(A) tail length for all MT-mRNA in 3T3 and ES measured by PAIso-seq2. Histograms (bin size = 1 nt) are normalized to cover the same area. Transcripts with poly(A) tail of at least 1 nt are included in the analysis. Transcripts with poly(A) tail length greater than 150 nt are included in the 150 nt bin. **e,** Box plot for the poly(A) tail length of each mitochondrial coding genes measured by PAIso-seq2. The “×” indicates the mean value, the horizontal bars show the median value, and the top and bottom of the box represent the values of the 25^th^ and 75^th^ percentiles, respectively. Transcripts with poly(A) tail of at least 1 nt for the given gene are included in the analysis. **f,** Proportion of MT-mRNA containing U, C, and G residues in overall, the 5’-end, internal, and 3’-end of the poly(A) tail in 3T3 and ES measured by PAIso-seq2. Transcripts with poly(A) tail of at least 1 nt are included in the analysis. Error bars indicate SEM from two replicates. **g,** Proportion of transcripts containing U (left), C (middle), and G (right) residues for each mitochondrial coding genes measured by PAIso-seq2. Transcripts with poly(A) tail of at least 1 nt are included in the analysis.

Poly(A) tails are important for the stability of nuclear transcripts, leading us to assess whether the poly(A) tail is also an essential posttranscriptional mechanism for regulating the stability of mitochondrial transcripts. We analyzed the distribution pattern of poly(A) tail lengths for all MT-mRNAs and each mitochondrial protein-coding gene. We found a similar distribution of poly(A) tail of all MT-mRNAs in 3T3 and ES (Fig. 1d) and found that the poly(A) tail length distributed in a narrow range (60 ± 30 nt) for all mitochondrial coding genes, except *Nd6* (Fig. 1e), which is known to have no poly(A) tail^12,20^.

Non-A residues can be incorporated in the poly(A) tail of nuclear transcripts and play key roles in RNA metabolism^18,21^. Therefore, we explored whether MT-mRNA contains non-A residues in the poly(A) tail. We found the presence of non-A residues in the 5’-end, internal, and 3’-end of the poly(A) tails of MT-mRNAs (Fig. 1f). In addition, the proportion of MT-mRNAs containing non-A residues differed between 3T3 and ES (Fig. 1f, g).

### MT-mRNA possesses a stable distribution pattern of poly(A) tail length

The poly(A) tail length of MT-mRNA is similar between 3T3 and ES, leading us to analyze whether it varies or stable among tissues. As such, we examined the mitochondrial transcriptome across 13 mouse tissues and found that the distribution pattern of the poly(A) tail length of MT-mRNA was parallel and stable in these tissues (Fig. 2a). Considering that the poly(A) tail length of nuclear transcripts is highly dynamic during the mammalian OET^22–26^, we analyzed whether this also happens to MT-mRNA. We found that MT-mRNA presented an extremely stable distribution pattern of its poly(A) tail length during oocyte maturation and preimplantation development in mice, rats, pigs, and humans. This was proven by both PAIso-seq1 and PAIso-seq2 (Fig. 2b-h). We also discovered that the distribution pattern of poly(A) tail length of MT-mRNA was similar across samples from these four species (Fig. 2b-h). These results indicate an unexpected feature of MT-mRNA that their poly(A) tail length follows a stable distribution pattern across diverse tissues and species.

**Fig. 2.**
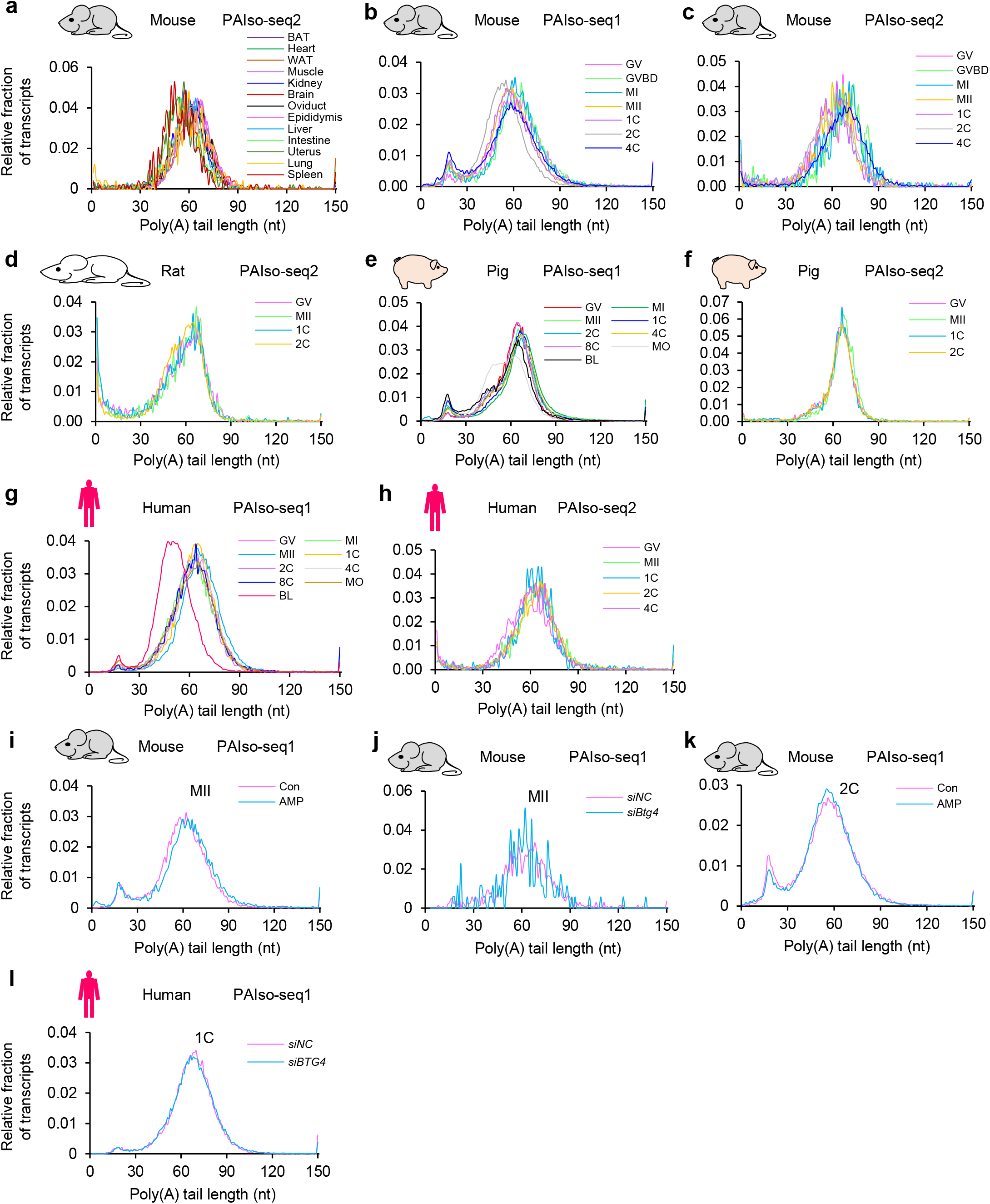
Highly consistent distribution pattern of MT-mRNA poly(A) tail length. **a,** Histogram of poly(A) tail length for all MT-mRNA across 13 mouse tissues measured by PAIso-seq2. BAT, brown fat; WAT, white fat. **b-h,** Histogram of poly(A) tail length for all MT-mRNA in oocytes and embryos at different stages in mice (**b**, **c**), rats (**d**), pigs (**e**, **f**), and humans (**g**, **h**) measured by PAIso-seq1 (**b**, **e**, **g**) or PAIso-seq2 (**c**, **d**, **f**, **h**). GV, germ-vesicle stage oocyte; MI, Metaphase I oocyte; MII, Metaphase II oocyte; 1C, 1-cell embryo; 2C, 2-cell embryo; 4C, 4-cell embryo; 8C, 8-cell embryo; MO, morula; BL, blastocyst. **i, k,** Histogram of poly(A) tail length for all MT-mRNA with or without addition of adenosine monophosphate (AMP) in mouse MII oocytes (**i**) or 2C embryos (**l**) measured by PAIso-seq1. **j, l**, Histogram of poly(A) tail length for all MT-mRNA with or without knockdown of *Btg4* in mouse MII oocytes (**j**) or human 1C embryos (**m**) measured by PAIso-seq1. In all panels, histograms (bin size = 1 nt) are normalized to cover the same area. Transcripts with poly(A) tail of at least 1 nt are included in the analysis. Transcripts with poly(A) tail length greater than 150 nt are included in the 150 nt bin.

The addition of adenosine monophosphate (AMP) can inhibit deadenylation of the poly(A) tail for nuclear transcripts by potently inhibiting the major deadenylase CNOT6L and CNOT7^27^, leading us to ask whether the poly(A) tail length of MT-mRNAs was affected by adding AMP. Our results demonstrated that AMP addition did not affect the distribution of MT-mRNA in the poly(A) tail length in both mouse metaphase II (MII) oocytes and mouse 2-cell embryos (Fig. 2i, k). It has been reported that BTG4 regulates the deadenylation and decay of thousands of maternal RNAs during the OET^23,24,28,29^. After assessing whether Btg4 regulates the deadenylation of MT-mRNA, we found no obvious change in the distribution pattern of poly(A) tail length of MT-mRNA upon Btg4 knockdown in either mouse MII oocytes or human 1-cell embryos (Fig. 2j, l). These results indicate that different regulation mechanisms are working on the poly(A) tail length dynamic between nuclear and mitochondrial mRNA transcripts.

### Non-A residues in MT-mRNA poly(A) tails vary across tissues and increase slightly during the OET

The poly(A) tail length distribution pattern of MT-mRNA is extremely stable across different tissues and developmental stages. As such, we analyzed the ratio of transcripts containing non-A residues to identify if they were stable across multiple tissues or during the developmental process. We calculated the proportion of MT-mRNA containing non-A residues across 13 mouse tissues and found slight differences among these tissues, particularly in the C residues (Fig. 3a). We next compared the proportion of MT-mRNA containing non-A residues during the OET in mice, rats, pigs, and humans. We detected a slight upward trend of the ratio of C residues in MT-mRNA, while no difference was found for U or G residues during the OET in mice and rats from both PAIso-seq1 and PAIso-seq2 (Fig. 3b-d). During the OET in pigs, the ratio of C and G residues increased (Fig. 3e, f), while during the OET in humans, the ratio of U, C, and G residues was slightly elevated after analysis with both PAIso-seq1 and PAIso-seq2 (Fig. 3g, h). These results indicate that the ratio of transcripts containing non-A residues of MT-mRNA varies slightly across mouse tissues, and increases slightly during OET in mice, rats, pigs, and humans.

**Fig. 3.**
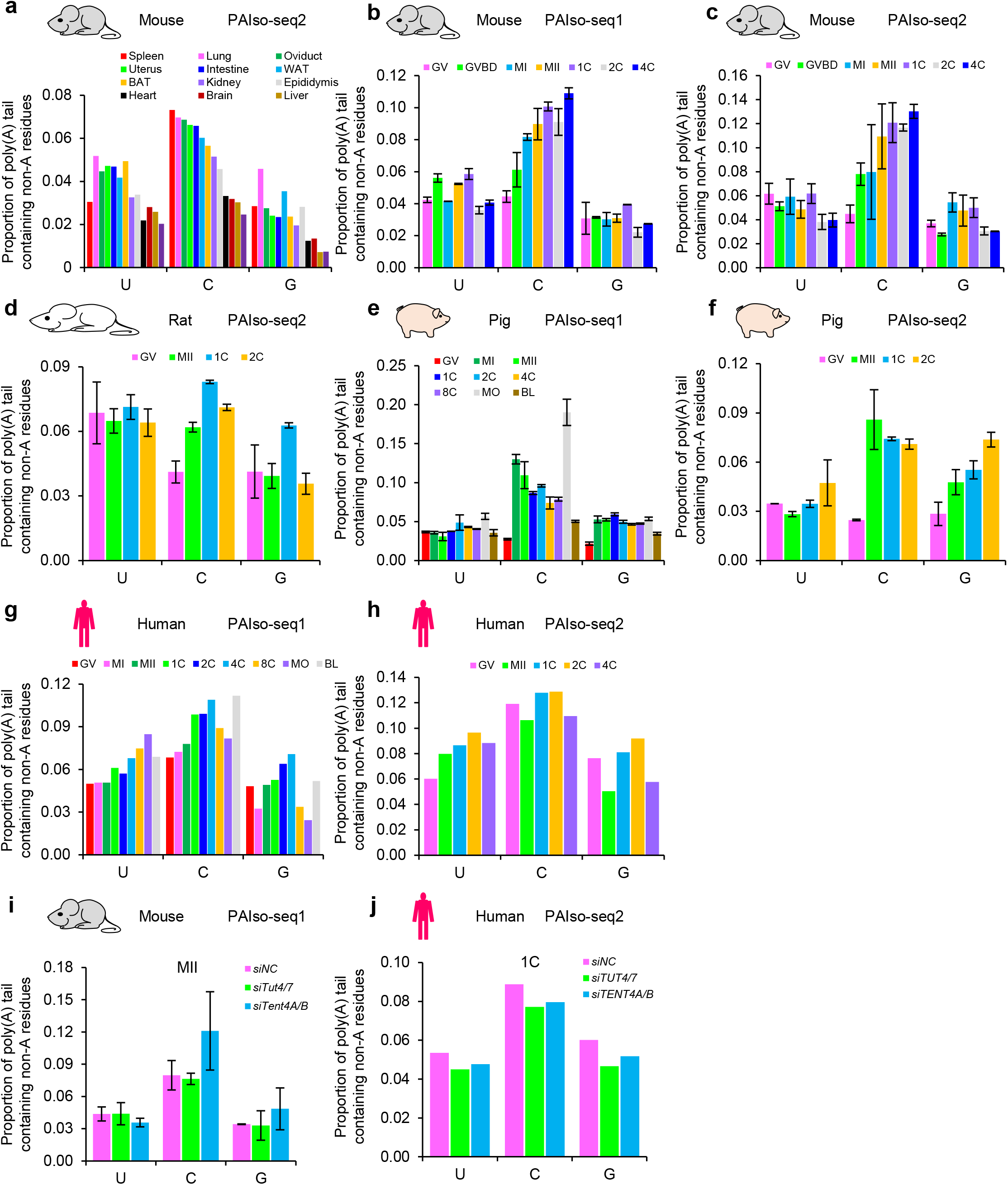
Slight variation of non-A residues in MT-mRNA poly(A) tails. **a,** Proportion of MT-mRNA containing U, C, and G residues across 13 mouse tissues measured by PAIso-seq2. Transcripts with poly(A) tail of at least 1 nt are included in the analysis. **b-h,** Proportion of MT-mRNA containing U, C, and G residues in oocytes and embryos at different stages in mice (**b**, **c**), rats (**d**), pigs (**e**, **f**), and humans (**g**, **h**) measured by PAIso-seq1 (**b**, **e**, **g**) and PAIso-seq2 (**c**, **d**, **f**, **h**). Transcripts with poly(A) tail of at least 1 nt are included in the analysis. Error bars indicate SEM from two replicates. **i, j,** Proportion of MT-mRNA containing U, C, G residues with or without knockdown of *Tut4/7* or *Tent4A/B* in mouse MII oocytes measured by PAIso-seq1 (**i**) or human 1C embryos measured by PAIso-seq2 (**j**). Transcripts with poly(A) tails of at least 1 nt are included in the analysis. Error bars indicate SEM from two replicates.

Considering that TUT4/7 and TENT4A/B incorporate non-A residues into the poly(A) tail of nuclear transcripts^23,25,30–34^, we assessed whether they are involved in non-A residues incorporation into poly(A) tail of MT-mRNA. We detected minimal changes in the ratio of transcripts containing non-A residues in MT-mRNA upon knockdown of either *Tut4/7* or *Tent4A/B* in mouse MII oocytes (Fig. 3i) and human 1-cell embryos (Fig. 3j), respectively. These results indicate that different regulation mechanisms are dictating how non-A residues are incorporated into the poly(A) tail between nuclear and mitochondrial mRNA.

### MT-mRNA abundance varies substantially across tissues and increases dramatically during the OET

The number of mitochondria in a cell can vary widely by tissue and cell type, leading us to study whether MT-mRNA abundance varies across multiple tissues or during the mammalian OET. We first examined MT-mRNA expression across 13 tissues in mice. We then determined the ratio of reads from mitochondria protein-coding genes to reads from nuclear protein-coding genes as mitochondrial mRNA abundance. As a result, we discovered that the abundance of mitochondrial mRNA is higher in tissues with high-energy demands, such as brown fat (BAT), heart, and muscle (Fig. 4a). This is consistent with results obtained by other researchers^12^. Since both the number of mitochondria and transcriptional activity contributes to mitochondrial mRNA abundance, this analysis provides an indirect assessment of the demand on mitochondria from different tissues.

**Fig. 4.**
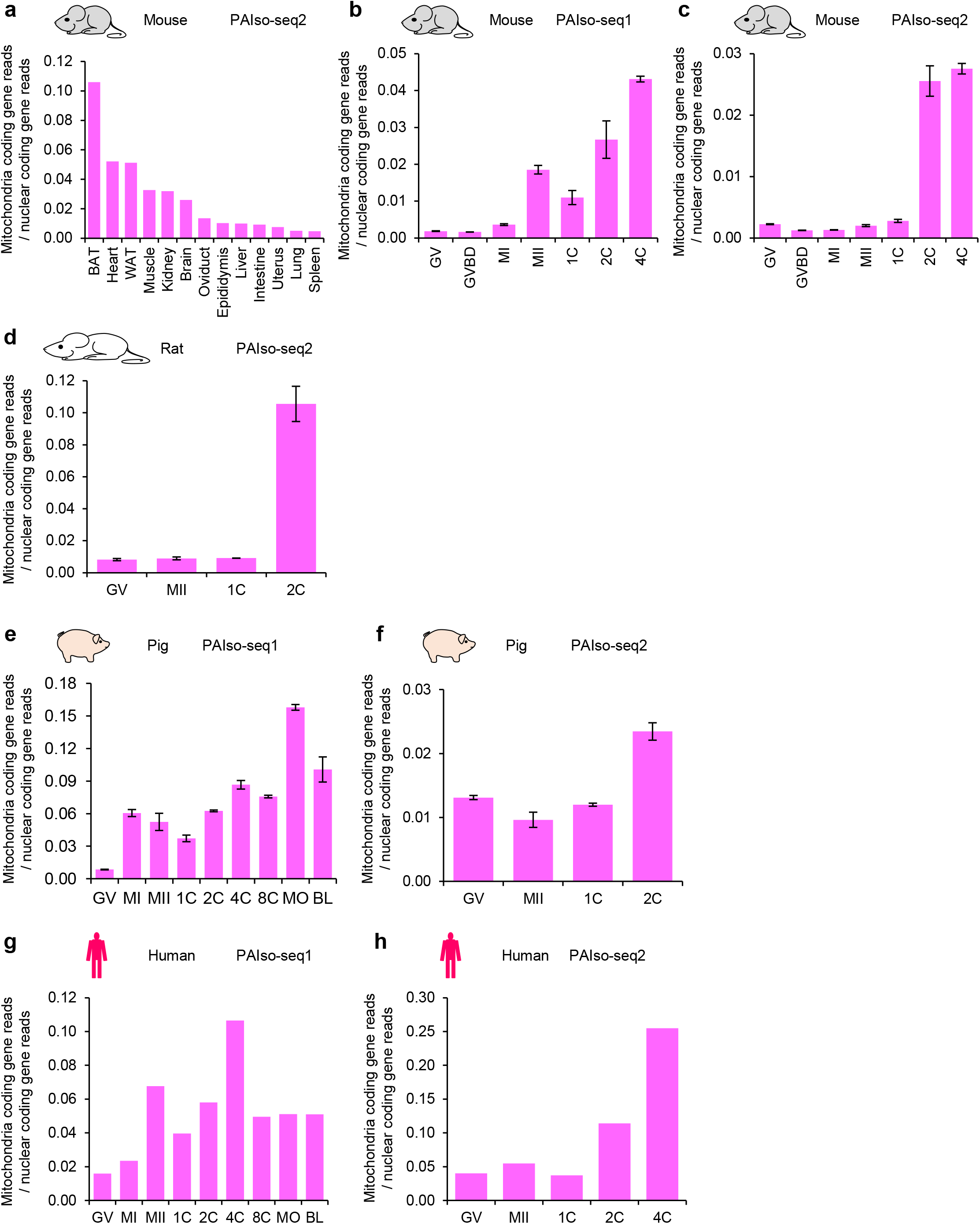
MT-mRNA abundance varies widely across tissues and elevates dramatically during mammalian OET. **a,** MT-mRNA abundance across 13 mouse tissues measured by PAIso-seq2. **b-h,** MT-mRNA abundance dynamic in oocytes and embryos at different stages in mice (**b**, **c**), rats (**d**), pigs (**e**, **f**), and humans (**g**, **h**) measured by PAIso-seq1 (**b**, **e**, **g**) or PAIso-seq2 (**c**, **d**, **f**, **h**). Error bars indicate SEM from two replicates.

We next examined MT-mRNA abundance during the OET in mice, rats, pigs, and humans and found dramatic increases in MT-mRNA abundance at the 2-cell stage in both mice and rats (Fig. 4b-d). This is typically when the major wave of zygotic genome activation (major ZGA) happens. The nuclear localization of mitochondrial TCA cycle enzymes is a critical step for mammalian major ZGA^9^, while our results indicated that increases of MT-mRNA abundance during the OET coincide with the occurrence of nuclear genome activation. Based on data from PAIso-seq1 and PAIso-seq2, we found elevated MT-mRNA abundance at the morula stage in pigs and at the 4-cell stage in humans (Fig. 4e-h), suggesting that the ZGA start around the morula stage in pigs and the 4-cell stage in humans.

Major ZGA is impaired in parthenogenetic embryos^26^. Therefore, we tested whether the abundance of the MT-mRNA was affected. We found that the MT-mRNA abundance was lower in parthenogenetic two-cell embryos than in fertilized two-cell embryos in both mice and rats (Fig. 5a, b). This result suggest that lower MT-mRNA abundances could be one reason for impaired ZGA in parthenogenetic embryos.

**Fig. 5.**
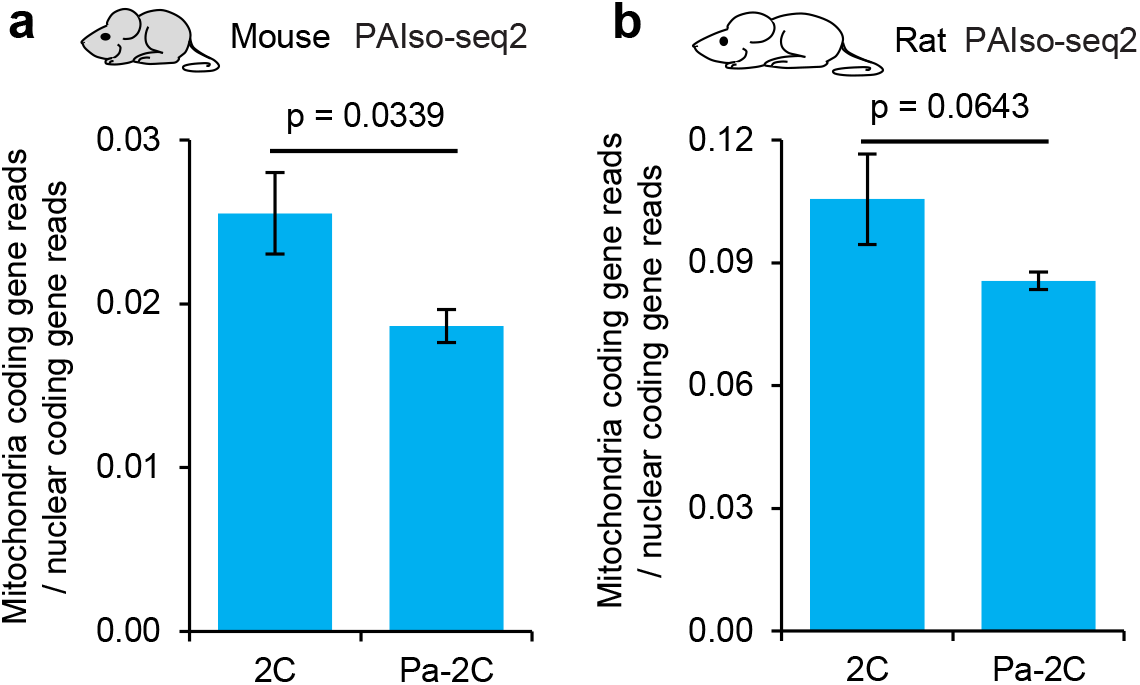
MT-mRNA abundance decreases in mouse and rat parthenogenetic embryos. **a-b,** MT-mRNA abundance in fertilized and parthenogenetic 2C (Pa-2C) embryos tin mice (**a**) and rats (**b**) measured by PAIso-seq2. Error bars indicate SEM from two replicates. *P* value is calculated by Student’s *t* test.

## Discussion

While mitochondria uniquely regulate their own gene expression, little is known about mitochondrial transcriptome features. In this study, we described these MT-mRNA features, including poly(A) tail length, non-A residues in the poly(A) tail, and abundance, in different cell lines, across multiple tissues, and during the mammalian OET. We discovered that the poly(A) tail length distribution pattern of MT-mRNA is extremely stable across cell lines, tissues and during mammalian oocyte maturation and preimplantation development (Fig. 6). We found that the ratio of poly(A) tails containing non-A residues and the abundance of mitochondria mRNA varies widely by cell lines and tissues (Fig. 6). Moreover, we found that the ratio of non-A residues of MT-mRNA slightly increases and that the abundance of MT-mRNA significantly increases along major ZGA during the OET in mice, rats, pigs, and humans (Fig. 6). This suggests a link between transcription of MT-mRNAs and nuclear genome transcriptional activation. However, whether there is a big wave of transcriptional activation of the mitogenome together with major ZGA of the nuclear genome during preimplantation development requires additional study.

**Fig. 6.**
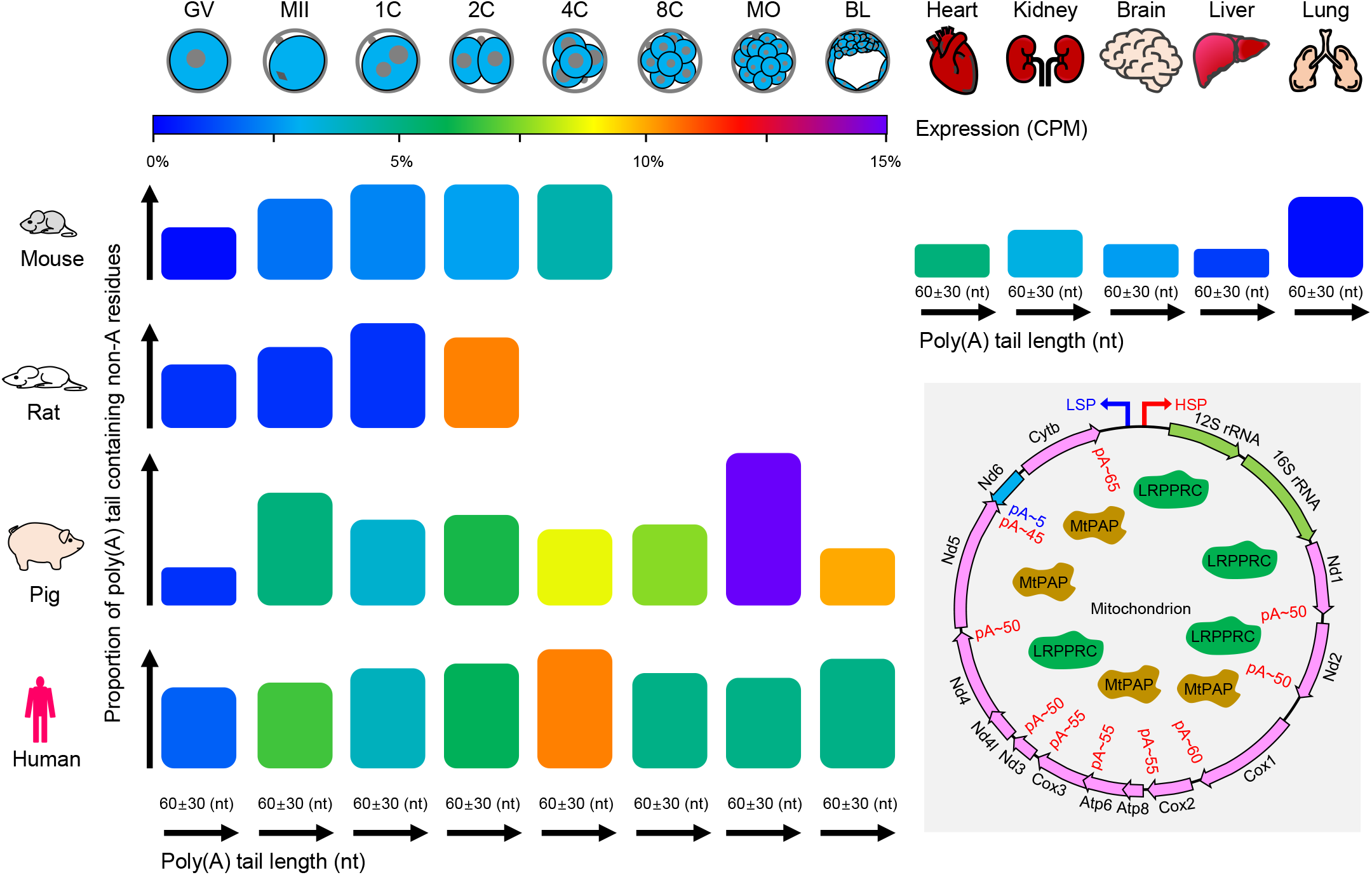
Model for dynamics of MT-mRNA features during the OET and across tissues. Dynamic of MT-mRNA features, including poly(A) tail length, ratio of tail non-A residues, and abundance, during the OET and across tissues. For MT-mRNA during OET in mice, rats, pigs, and humans, poly(A) tail length distribution is extremely stable, the ratio of non-A residues elevates slightly, and the abundance significantly increases. For MT-mRNA across mouse tissues, the poly(A) tail length distribution is stable, and the ratio of non-A residues and abundance varies widely. The median poly(A) length for each mitochondrial coding genes is around 45-65 nt, except *Nd6* which is only about 5 nt. Enzymes that could be involved in poly(A) tail processing are also shown in the model, such as MtPAP (mitochondrial poly(A) polymerase) and LRPPRC (leucine-rich PPR-motif containing).

Translational regulation of mRNA through poly(A) tail length have been reported in neurons, immune cells and oocytes^24,35^. Surprisingly, the poly(A) tail of MT-mRNAs maintain similar length across cell types, developmental stages, and species. In particular, although the global poly(A) tail length of mRNA from nuclear protein-coding genes changes substantially between stages during OET in mice, rats, pigs and humans^22–26^, the poly(A) tail length of MT-mRNAs are unchanged during OET in these four species. Therefore, it is likely that the translation of MT-mRNAs are not actively regulated by the poly(A) tail length. In addition, the *Nd6* is generally not polyadenylated, the translation of which does not require poly(A) tails, suggesting a unique mechanism on regulating the stability and translation of *Nd6*.

Mitochondria are semi-autonomous organelles. We discovered that the poly(A) tailing mechanism on MT-mRNA differs from that of nuclear mRNA. It is unclear whether other post-transcriptional modifications, including chemical modifications such as N^6^-methyl-adenosine (m6A) and N4-acetylcytidine (ac4C), are present in MT-mRNA and added or removed from the MT-mRNA by the same mechanisms as those in the nuclear mRNA. Chloroplasts are another kind of semi-autonomous organelle; it will be interesting to ask whether the post-transcriptional modifications of mRNA in chloroplasts are conserved with mitochondrial or nuclear mRNA. These questions warrant further study.

## Material and Methods

### Poly(A) analysis for mitochondrial mRNA

PAIso-seq1 and PAIso-seq2 sequencing data processing are described in our previous work (our mouse/human paper). Clean CCS reads were aligned to Mus musculus UCSCmm10 reference genome using minimap2 (v.217-r941). The annotation information of the mitochondrial coding gene *Atp8* and *Nd4l* were deleted and the names of both *Atp6* and Nd4 were changed to *Atp6/8* and *Nd4/4l* in the genome annotation files (.gtf) of mice, rats, pigs, and humans. This produced new genome annotation files, which were used to annotate the genes. The poly(A) tail was then extracted using python script PolyA_trim.py (http://). The 3’-soft clip sequence of the CCS reads in the alignment file is used as a candidate poly(A) tail sequence. The number of A, U, C, and G residues were counted in the 3’-soft clip sequences of each alignment. The 3’-soft clip sequences with the frequency of U, C, and G greater or equal to 0.1 simultaneously were marked as “HIGH_TCG” tail. To better definite the poly(A) tail, we defined a continuous score based on the transitions between the two adjacent nucleotide residues throughout the 3’-soft clip sequences. To calculate the continuous score, the transition from one residue to the same residue was scored as 0, and the transition from one residue to a different residue was scored as 1. The 3’-soft clips that were not marked as “HIGH_TCG” and which had a continuous score less than or equal to 12 were considered to be a poly(A) tail. Finally, we analyzed the base composition of the poly(A) tail using python script PolyA_note.py (http://).

### Statistical analyses

Statistical analyses [mean ± standard error of the mean (SEM)] were performed in Excel. Levels of significance were calculated using Student’s *t*-tests.

### Data availability

The sequencing data of PAIso-seq2 on 3T3 cells, mES cells, 13 mouse tissues (uterus, liver, brain, brown fat, lung, kidney, spleen, oviduct, intestine, epididymis white fat, and muscle), mouse oocytes (GV, GVBD, MI, and MII stages), mouse embryos (1C, 2C, and 4C stages, and pathogenetic 2C), rat oocytes (GV and MII stages), rat embryos (1C and 2C stages, and pathogenetic 2C), pig oocytes (GV and MII stages), pig embryos (1C and 2C stages), human oocytes (GV and MII stages), human embryos (1C, 2C, and 4C stages), and the sequencing data of PAIso-seq1 on mouse oocytes (GV, GVBD, MI, and MII stages, AMP treated MII, and *Btg4* knockdown MII), mouse embryos (1C, 2C, and 4C stages, and AMP treated 2C), pig oocytes (GV, MI, and MII stages), pig embryos (1C, 2C, 4C, 8C, MO, and BL stages), human oocytes (GV, MI, and MII stages), human embryos (1C, 2C, 4C, 8C, MO, and BL stages, and *BTG4* knockdown 1C), have been described in other papers (our mouse/human/pig paper). The ccs data in bam format from PAIso-seq1 and PAIso-seq2 experiments will be available at Genome Sequence Archive hosted by National Genomic Data Center. Custom scripts used for data analysis will be available upon request.

## Acknowledgments

We thank Hu Nie for his technical assistance in bioinformatic analysis. This work was supported by the National Key Research and Development Program of China (2018YFA0107001, 2016YFA0100200), the Strategic Priority Research Program of the Chinese Academy of Sciences (XDA24020203), National Natural Science Foundation of China (31970588, 32170606), Natural Science Foundation of Heilongjiang province (YQ2020C003), the China Postdoctoral Science Foundation (2020M670516, 2020T130687), the State Key Laboratory of Molecular Developmental Biology, and the Heilongjiang Touyan Innovation Team Program.

## Author contributions

Yusheng Liu, Falong Lu and Jiaqiang Wang conceived the project and designed the study. Yusheng Liu, Yiwei Zhang, Falong Lu and Jiaqiang Wang analyzed the sequencing data. Zhonghua Liu provided funding support for the project. Yusheng Liu and Jiaqiang Wang organized all figures. Yusheng Liu, Falong Lu and Jiaqiang Wang supervised the project. Yusheng Liu, Falong Lu and Jiaqiang Wang wrote the manuscript with the input from the other authors.

